# Exploring the Stony Coral Tissue Loss Disease Bacterial Pathobiome

**DOI:** 10.1101/2020.05.27.120469

**Authors:** D.D. Iwanowicz, W.B. Schill, C. M. Woodley, A. Bruckner, K. Neely, K.M. Briggs

## Abstract

A devastating novel coral disease outbreak, referred to as Stony Coral Tissue Loss Disease (SCTLD), was first described in 2014. It is thought to have originated offshore of Miami-Dade County, FL, but has persisted and spread, affecting new reefs along the Florida Reef Tract and reefs of at least 8 other Caribbean jurisdictions. We investigated the microbial communities of clinically normal and diseased specimens of five species of affected corals using targeted 16S ribosomal DNA sequencing (Illumina MiSeq). Fifty-nine bacterial sequences were identified using contrast analysis that had enriched abundance in diseased coral host microbiomes relative to the microbiomes of clinically normal hosts. Several sequences from known bacterial pathogens were identified in this group. Additionally, we identified fifty-three bacterial species that had differentially elevated numbers in clinically normal coral host samples relative to samples from diseased host corals. The bacterial consortia composing the clinically normal and diseased coral microbiomes were clearly distinct taxonomically. Predicted functional profiles based on taxonomy, however, were found to be quite similar. This indicates a high level of functional redundancy among diseased and clinically normal microbiome members. Further examination of the direct sequencing data revealed that while some bacteria were differentially distributed according to disease status, others were not. Fifty-one bacterial species were found in both diseased and clinically normal coral host samples and not differentially abundant in either disease state. These still may be important in explaining the presentation of disease.

**IMPORTANCE:** Determining causation is a management top priority to guide control and intervention strategies for the SCTLD outbreak. Towards this goal we examined bacterial taxa that were differentially elevated in numbers in diseased corals as compared to clinically normal corals at Looe Key, FL in August 2018. Many of the bacterial species we detected are known to be pathogenic to humans, animals, and (or) plants, and some of these have been found associated with diseased corals in other studies. Microbes that were present (or conspicuous by their absence) in both diseased as well as clinically normal corals were also examined because “healthy” corals from a diseased location such as Looe Key may have been exposed but may not have been showing overt disease at the time of sampling. Although untangling of causation is not possible currently, certain bacterial cliques and excess nutrients appear to be potential risk factors in SCTLD pathology.

---

Stony coral tissue loss disease (SCTLD) is a newly emerged lethal disease that has been affecting scleractinian corals along the Florida Reef Tract (FRT) since 2014. With white plague-like signs (https://nmsfloridakeys.blob.core.windows.net/floridakeys-prod/media/docs/20181002-stony-coral-tissue-loss-disease-case-definition.pdf), this disease outbreak was first recorded at Virginia Key near Miami (1, 2). By the winter of 2015, the disease had spread to the northern area of the Upper Keys within the Florida Keys National Marine Sanctuary. In 2016 and 2017 the disease spread north through Martin County and south through the Upper Keys (2). Between 2014-2015, SCTLD was found by W. F. Precht et al. (1) to spread at a rate of 2.5-5 km per month, but this rate of disease advancement apparently increased to 8-22 km per month in 2017-2018 as noted by K. Neely (3). In 2018, researchers were observing corals infected with SCTLD in the Middle Keys, and by December of 2019 corals infected by SCTLD were found well south of Key West and into the Marquesas Keys. Though originating in Florida, signs of SCTLD were recognized in Jamaica in 2017 and the disease now has been reportedly observed or suspected in the Mexican Caribbean, St. Maarten, the U.S. Virgin Islands, the Dominican Republic, the Turks and Caicos, Belize, the Netherlands Antilles, Puerto Rico, and the Bahamas (2).

SCTLD has been found to affect more than 20 of the 45 Caribbean reef-building coral species (stony corals), does not seem to be self-limiting as in the case of some other diseases, and does not seem to slow in cooler seasons (2). The affected species include four of seven Caribbean species that are listed as threatened (Federal Register 2006, 2014) under the Endangered Species Act of 1973 and have been further categorized as either highly susceptible, intermediately susceptible, presumed susceptible, and low susceptible species. The disease causes rapid mortality among affected coral species, with a high rate of transmission and it has affected a large geographic range over the more than six years it has been spreading (2). It is interesting to note that neither *Acropora palmata* nor *A. cervicornis* show susceptibility to SCTLD. As most reported coral diseases have low prevalence (<5%) and are not considered contagious (1), this multi-year outbreak is somewhat of a unique phenomenon. The gross morphological characteristics of the disease can vary among sites and species. Tissue loss often appears basally, peripherally or both on a colony and spreads upward leaving intact white skeleton (indicative of the rapid tissue loss) before algal colonization follows within 3-7 days. Tissue loss lesions can also present as patches within intact tissue that increase in size, often fusing into larger denuded areas. Histologically, the lesions appear first along the basal body wall then progress upward to envelop the rest of the polyp (J. H. Landsberg, unpublished data; https://nmsfloridakeys.blob.core.windows.net/floridakeys-prod/media/docs/20181002-stony-coral-tissue-loss-disease-case-definition.pdf).

Causative agents for coral reef declines appear to be diverse and complex. Risk factors include water pollution, habitat destruction, overfishing, invasive species, global climate change, and coral diseases (4, 5). This outbreak has precipitated increased monitoring and research efforts on SCTLD including its etiology, and transmission dynamics, along with intervention actions to halt or minimize impacts. Interventions including treatment of diseased colonies with specific antibiotics have been shown to retard or stop the tissue loss suggesting bacterial involvement in the disease etiology (6–11). The similar disease presentation and response to treatment across many coral species suggests that the same causative agent(s) are at work with limited variation among coral hosts.

Metagenomic sequencing of the 16S rRNA gene has been widely used to study the bacterial composition of multiple and varied sample types, most notably perhaps, the human microbiome (12–15). Increasingly however, metagenomic studies have been made on an ever broadening array of environmental microbial consortia including those associated with plants, animals, water, and sediment (16–19) as well as those associated with corals (20–25).

The aim of this study was to identify possible causative agents for SCTLD that have been affecting the Florida Reef Tract since 2014. We report here the bacterial consortia associated with five SCTLD-susceptible coral species (FIG 1) collected from Looe Key, FL in 2018 during an active outbreak. Direct, high throughput 16S amplicon sequencing (Illumina MiSeq) was used to determine if or how microbial communities differ between clinically normal and diseased corals.

**FIG 1.**
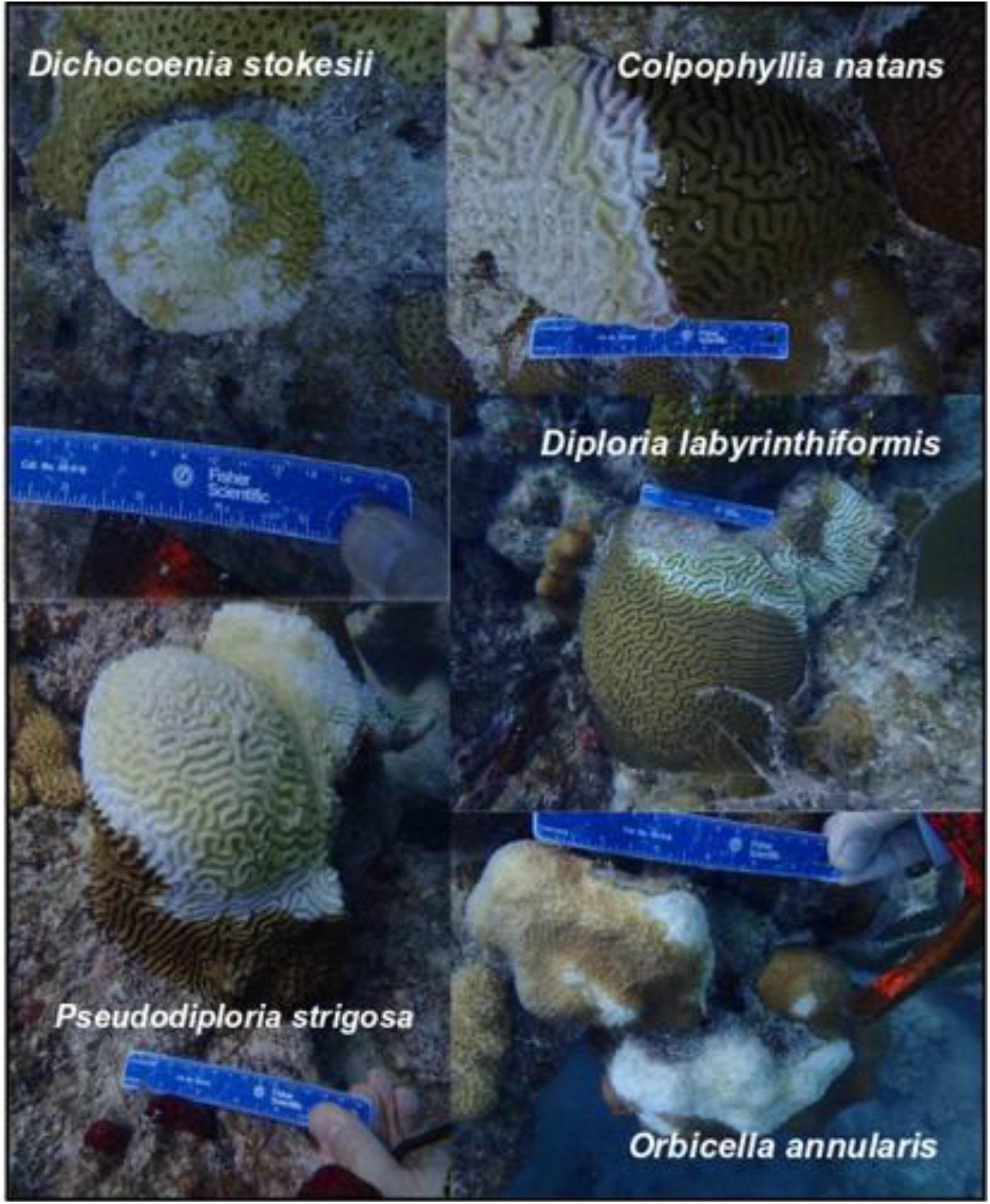
Representative samples of five coral species affected by Stony Coral Tissue Loss Disease (SCTLD) at Looe Key, FL in August 2018.

## RESULTS

We performed 16S sequencing of the bacterial consortia associated with diseased and clinically normal specimens of five important species of Caribbean stony corals affected by SCTLD to identify potential causative agents. A total of 10,081,496 sequence reads passed quality filtering with 6,410,207 of these classified by the One Codex platform at the species level and 1,546 total bacterial species were identified in the diseased coral samples. Bacterial species detected in the clinically normal coral samples totaled 1,981. After trimming uninformative species classifications with the lowest read assignments (lower 0.25%), 1,119 and 1,407 species classifications remained from the diseased and clinically normal samples, respectively. Remaining sequence assignments for all libraries numbered minimally 96 to 1,961 depending on the host sample (Table 2). Thus, the classifications retained for further analyses were those that were observed a reasonable number of times.

**Table 1.** Site number, latitude, longitude, Scientific Name, Abbreviation, Common Name, Status (CN = Clinically Normal, D = Diseased), Depth of Collection (meters), and sample identification of 19 corals collected from Looe Key, Florida.

| Site | Latitude | Longitude | Scientific Name | Abbreviation | Common Name | Status | Depth (m) | Sample ID |
| --- | --- | --- | --- | --- | --- | --- | --- | --- |
| 1 | N 24° 32' 40.812" | W 81° 24' 30.96" | *Colpophyllia natans* | CNAT | boulder brain coral | D | 6.71 | CNAT_DisBgF |
| 2 | N 24° 32' 44.988" | W 81° 24' 15.84" |  |  |  | D | 7.01 | CNAT_DisBgBBUnl |
| 2 | N 24° 32' 44.988" | W 81° 24' 15.84" |  |  |  | CN | 6.40 | CNATHC-8 |
| 2 | N 24° 32' 44.988" | W 81° 24' 15.84" |  |  |  | CN | 5.18 | CNATHC-9 |
| 1 | N 24° 32' 40.812" | W 81° 24' 30.96" | *Dichocoenia stokesii* | DSTO | elliptical star coral | D | 5.49 | DSTO_DisBgJ |
| 1 | N 24° 32' 40.812" | W 81° 24' 30.96" |  |  |  | D | 5.49 | DSTO_DisBgL |
| 1 | N 24° 32' 40.812" | W 81° 24' 30.96" |  |  |  | CN | 5.49 | DSTOHC-1 |
| 1 | N 24° 32' 40.812" | W 81° 24' 30.96" |  |  |  | CN | 5.49 | DSTOHC-3 |
| 1 | N 24° 32' 40.812" | W 81° 24' 30.96" | *Diploria labyrinthiformis* | DLAB | grooved brain coral | D | 6.40 | DLAB_DisBgA |
| 2 | N 24° 32' 44.988" | W 81° 24' 15.84" |  |  |  | D | 5.79 | DLAB_DisBgE |
| 2 | N 24° 32' 44.988" | W 81° 24' 15.84" |  |  |  | CN | 7.01 | DLABHC-5 |
| 1 | N 24° 32' 40.812" | W 81° 24' 30.96" | *Orbicella annularis* | OANN | lobed star coral | D | 7.01 | OANN_DisUNLB |
| 1 | N 24° 32' 40.812" | W 81° 24' 30.96" |  |  |  | D | 7.01 | OANN_DisBgD |
| 2 | N 24° 32' 44.988" | W 81° 24' 15.84" |  |  |  | CN | 5.49 | OANNHC-6 |
| 2 | N 24° 32' 44.988" | W 81° 24' 15.84" |  |  |  | CN | 5.79 | OANNHC-7 |
| 1 | N 24° 32' 40.812" | W 81° 24' 30.96" | *Pseudodiploria strigosa* | PSTR | symmetrical brain coral | D | 5.49 | PSTR_DisBgH |
| 1 | N 24° 32' 40.812" | W 81° 24' 30.96" |  |  |  | D | 5.18 | PSTR_DisBgI |
| 1 | N 24° 32' 40.812" | W 81° 24' 30.96" |  |  |  | CN | 5.49 | PSTRHC-2 |
| 1 | N 24° 32' 40.812" | W 81° 24' 30.96" |  |  |  | CN | 5.49 | PSTRHC-4 |

**Table 2.**
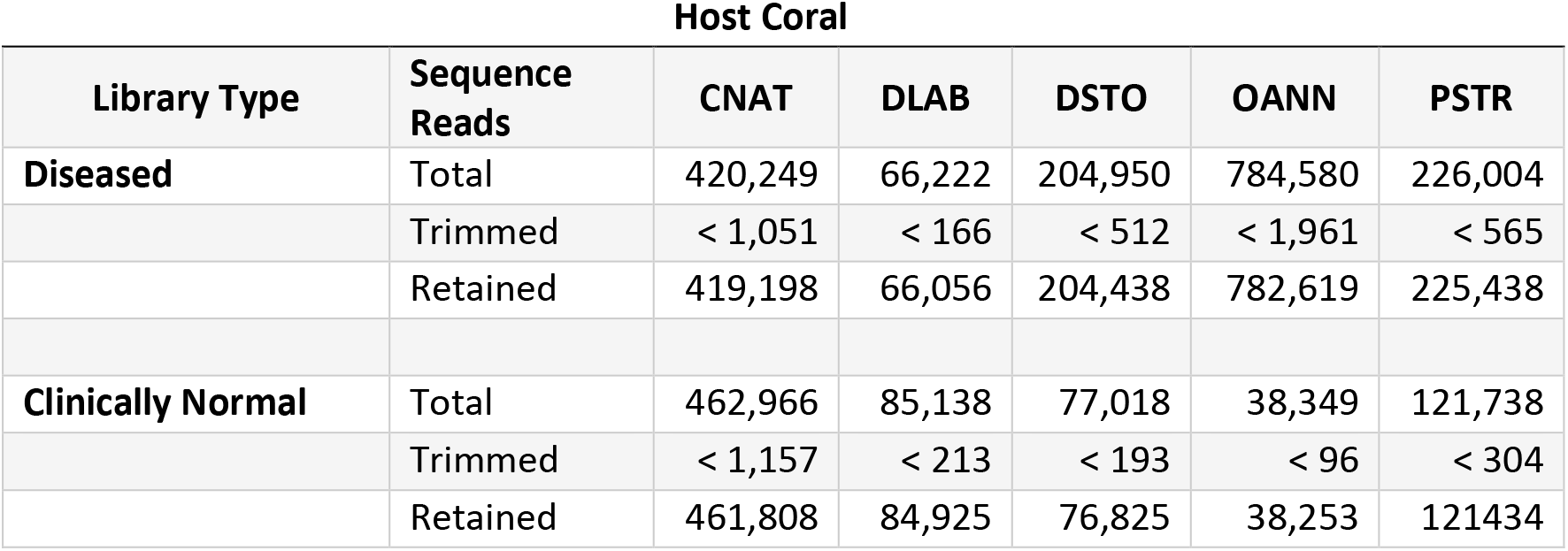
Total numbers of classified reads for each library type, reads trimmed (0.25% of total bacterial reads classified for each coral host species), and classified reads retained for analysis. CNAT = *Colpophyllia natans*: DLAB = *Diploria labyrinthiformis*; DSTO = *Dichocoenia stokesii*; OANN = *Orbicella annularis*; and PSTR = *Pseudodiploria strigosa*.

### Diversity analyses

We examined the distribution of genetic diversity within samples (alpha diversity) as well as among disease states (beta diversity). Alpha diversity measures (Simpson; FIG 2) were somewhat variable for some species and (or) disease state replicates, but these variations were not statistically significant among the coral host microbiomes that were tested regardless of host species and (or) disease status. The Mann-Whitney test yielded a p-value of 0.84211 for the comparison of diseased versus clinically normal alpha diversity values, and evaluation of differences in alpha diversity among coral species sampled using Kruskal-Wallis testing yielded a p-value of 0.50768. In contrast, beta diversity analysis of Jenson-Shannon divergence measures using ANOSIM demonstrated that the microbes associated with the diseased and clinically normal states were of quite different composition (FIG 3).

**FIG 2.**
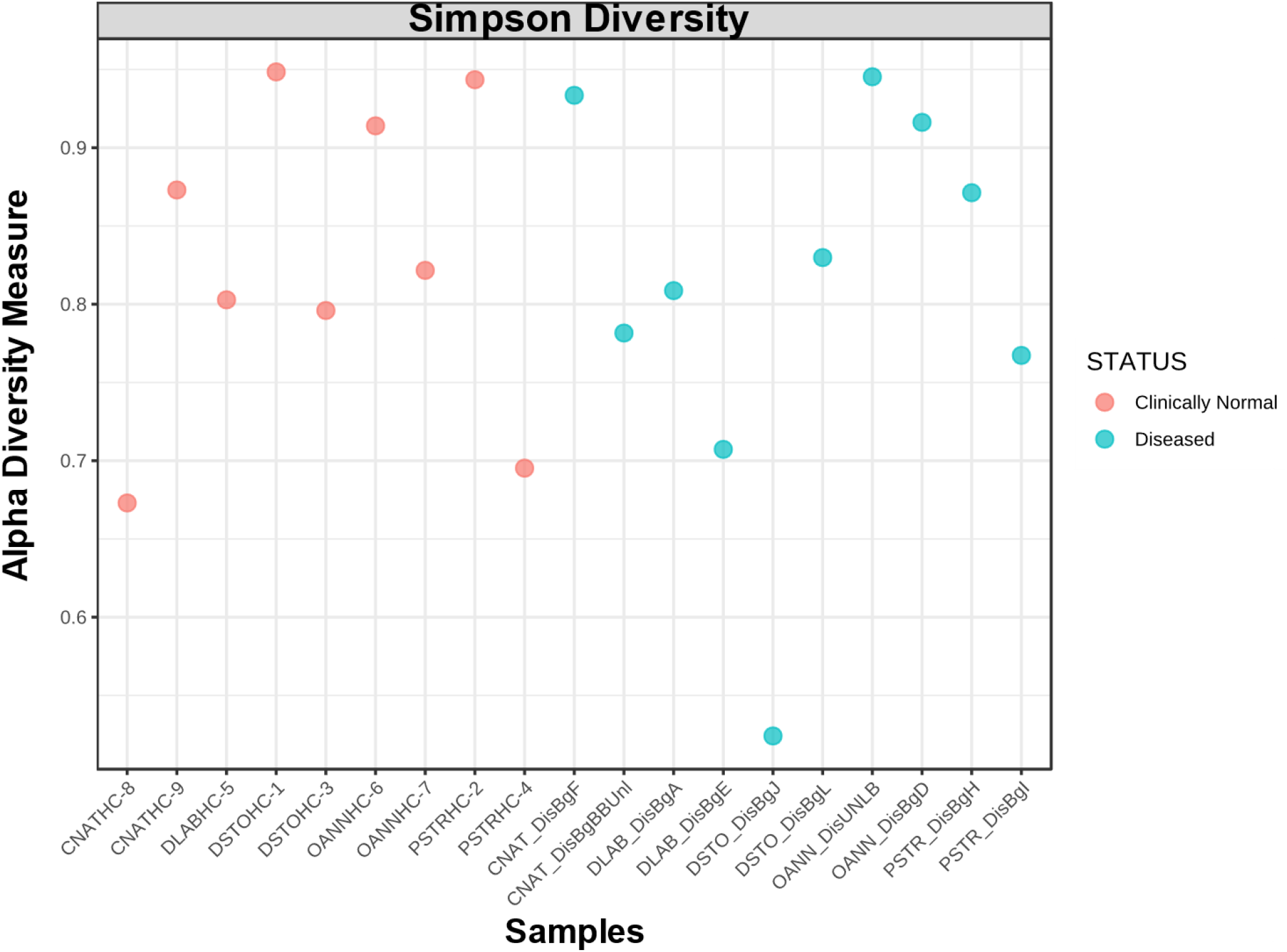
Alpha diversity as measured by the Simpson index was generally similar among the microbiomes of all five coral hosts species regardless of health status.

**FIG 3.**
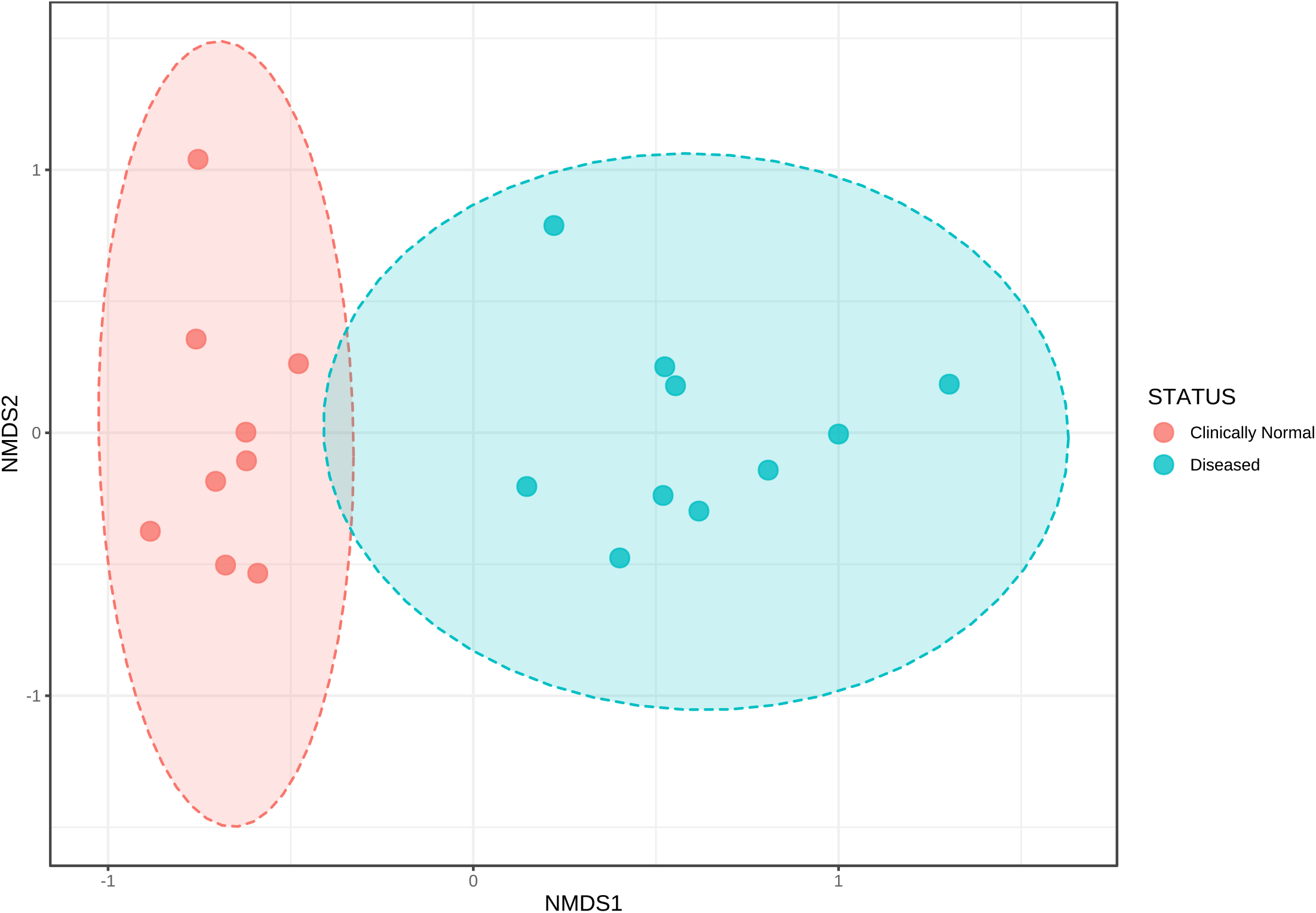
Nonmetric multidimensional scaling visualization of Jenson-Shannon Divergence measures using ANOSIM demonstrated that the microbiome taxa of five stony coral species affected by SCTLD are distinct between clinically normal and diseased colonies.

### Distribution of bacterial taxa across coral species and according to disease state

The distribution across host corals of the most commonly identified bacterial species are shown in Tables 3 and 4. Two species of *Achromobacter* and three species of *Fulvivirga* were common among the clinically normal coral microbiomes (Table 3) and *Achromobacter xylosoxidans* was universally present. Two species of Arcobacter were common among the diseased coral microbiomes with *Arcobacter bivalviorum* being a member of all diseased microbiome libraries as were *Algicola bacteriolytica* and *Clostridioides difficile.* Contrast analysis using edgeR revealed fifty-nine bacterial species that were enriched in abundance in diseased coral host samples relative to clinically normal hosts and are shown in Table 5. Fifty-three bacterial species had differentially elevated numbers in clinically normal coral host samples relative to diseased host corals (Table 6), while fifty-one bacterial species were found in both diseased and clinically normal coral host samples and not differentially abundant in either disease state (Table 7). All the taxa listed in Table 3 were found to be present in clinically normal as well as diseased samples (Table 7) except for *Fulvivirga kasyanovil*, found only in clinically normal samples (Table 6). All the taxa listed in Table 4 were found to be among those elevated in diseased coral samples (Table 5) except for *Arcobacter sp.* UDC415 and “*Candidatus* Amoebophilus asiaticus” that were present in both clinically normal and diseased coral samples (Table 7). Some bacteria known from other studies to be generally abundant and associated with a wide variety of corals and other marine invertebrates (22) were rare in the Looe Key coral microbiomes. *Ruegeria spp.*, for example, were relatively low in abundance and restricted to the clinically normal samples (Table 6). *Endozoicomonas* species abundances in clinically normal coral samples were found to be 0% in *C. natans* and *D. labyrinthiformis*, 0.186% in *D. stokesii*, 0.013% in *O. annularis*, and 1.604% in *P. strigosa*. *Endozoicomonas* in diseased coral samples was only detected in *O. annularis* at an abundance of 0.582%.

**Table 3.** Distribution of classified bacterial sequences across clinically normal host coral species. CNAT = *Colpophyllia natans*: DLAB = *Diploria labyrinthiformis*; DSTO = *Dichocoenia stokesii*; OANN = *Orbicella annularis*; and PSTR = *Pseudodiploria strigosa*.

| Microbe | CNAT | DLAB | DSTO | OANN | PSTR | Species Affected |
| --- | --- | --- | --- | --- | --- | --- |
| *Achromobacter xylosoxidans* | X | X | X | X | X | 5 |
| *Achromobacter denitrificans* | X | X |  | X | X | 4 |
| *"Candidatus Amoebophilus asiaticus"* | X | X |  | X | X | 4 |
| *Fulvivirga imtechensis* | X | X |  | X | X | 4 |
| *Fulvivirga kasyanovii* | X | X |  | X | X | 4 |
| *Fabibacter misakiensis* |  | X |  | X | X | 3 |
| *Fulvivirga lutimaris* | X | X |  | X |  | 3 |
| *Nitrincola lacisaponensis* | X |  | X |  | X | 3 |
| *Rhodospirillum rubrum* | X | X |  | X |  | 3 |
| *Sinorhizobium fredii* | X |  | X |  | X | 3 |
| 171 More- found on one or two hosts |  |  |  |  |  |  |

**Table 4.** Distribution of classified bacterial sequences across diseased coral hosts. CNAT = *Colpophyllia natans*: DLAB = *Diploria labyrinthiformis*; DSTO = *Dichocoenia stokesii*; OANN = *Orbicella annularis*; and PSTR = *Pseudodiploria strigosa*.

| Microbe | CNAT | DLAB | DSTO | OANN | PSTR | Species Affected |
| --- | --- | --- | --- | --- | --- | --- |
| *Algicola bacteriolytica* | X | X | X | X | X | 5 |
| *Arcobacter bivalviorum* | X | X | X | X | X | 5 |
| *Clostridioides difficile* | X | X | X | X | X | 5 |
| *Arcobacter* sp. UDC415 |  | X | X | X | X | 4 |
| "*Candidatus* Amoebophilus asiaticus" | X | X | X | X |  | 4 |
| *Pseudofulvibacter geojedonensis* |  | X | X |  | X | 3 |
| *Romboutsia lituseburensis* | X |  | X |  | X | 3 |
| *Shimia aquaeponti* | X |  | X | X |  | 3 |
| *Vibrio* sp. r24 | X |  | X |  | X | 3 |
| 95 More- found on one or two hosts |  |  |  |  |  |  |

**Table 5.** Bacterial species enriched in diseased coral hosts as determined by edgeR contrast analysis. Designations are predicated at a false discovery rate (FDR) < 0.01. Shown are the logarithm of 2-fold change (log2FC), the logarithm of counts per million reads (log_2_CPM), the Benjamini-Hochberg (B-H) adjusted P value (Pvalue), and the FDR. The table is sorted from highest to lowest abundance (log_2_CPM).

| Bacterial Species | log2FC | Log2CPM | Pvalue | FDR |
| --- | --- | --- | --- | --- |
| *Clostridioides difficile* | 5.832 | 16.425 | 3.160E-06 | 9.071E-06 |
| *Arcobacter bivalviorum* | 11.320 | 16.378 | 1.253E-11 | 2.042E-09 |
| *Tepidibacter mesophilus* | 8.471 | 13.556 | 1.669E-06 | 5.440E-06 |
| *Algicola bacteriolytica* | 4.113 | 13.416 | 1.801E-03 | 2.718E-03 |
| *Shimia aquaeponti* | 8.262 | 13.341 | 1.943E-07 | 1.131E-06 |
| *Vibrio* sp. r24 | 8.190 | 13.263 | 1.752E-08 | 2.561E-07 |
| *Marinilabilia nitratreducens* | 7.999 | 13.082 | 3.786E-07 | 1.870E-06 |
| *Burkholderia gladioli* | 7.912 | 12.996 | 4.535E-07 | 2.174E-06 |
| *Pseudofulvibacter geojedonensis* | 7.795 | 12.885 | 2.781E-06 | 8.243E-06 |
| *Marinifilum fragile* | 7.756 | 12.845 | 1.299E-06 | 4.504E-06 |
| *Staphylococcus epidermidis* | 3.957 | 12.802 | 4.630E-03 | 6.799E-03 |
| *Vibrio nigripulchritudo* | 7.559 | 12.638 | 3.033E-08 | 3.166E-07 |
| *Pseudahrensia aquimaris* | 7.477 | 12.577 | 1.191E-05 | 2.520E-05 |
| *Tenacibaculum litopenaei* | 7.367 | 12.464 | 2.712E-06 | 8.185E-06 |
| *Abyssivirga alkaniphila* | 4.250 | 12.215 | 2.063E-03 | 3.084E-03 |
| *Amphritea japonica* | 7.080 | 12.169 | 5.726E-08 | 5.490E-07 |
| *Romboutsia lituseburensis* | 7.009 | 12.094 | 1.885E-08 | 2.561E-07 |
| *Vibrio alginolyticus* | 6.990 | 12.070 | 2.569E-09 | 6.978E-08 |
| *Carboxylicivirga taeanensis* | 6.870 | 11.979 | 6.971E-06 | 1.623E-05 |
| *Meridianimaribacter flavus* | 6.601 | 11.713 | 1.571E-06 | 5.226E-06 |
| *Rhizobium* sp. Trp 3 | 6.397 | 11.519 | 4.939E-06 | 1.258E-05 |
| *Vibrio harveyi* | 6.314 | 11.409 | 2.097E-09 | 6.836E-08 |
| *Arcobacter molluscorum* | 6.236 | 11.364 | 4.615E-06 | 1.194E-05 |
| *Mesorhizobium* sp. | 6.210 | 11.341 | 9.035E-06 | 2.045E-05 |
| *Yersinia enterocolitica* | 6.165 | 11.286 | 5.160E-07 | 2.336E-06 |
| *Shimia isopora* | 6.062 | 11.193 | 1.990E-06 | 6.360E-06 |
| *Natronoflexus pectinivorans* | 5.979 | 11.117 | 6.027E-06 | 1.489E-05 |
| *Vibrio parahaemolyticus* | 5.961 | 11.072 | 4.088E-09 | 9.519E-08 |
| *Fusibacter paucivorans* | 5.876 | 11.019 | 4.533E-06 | 1.192E-05 |
| *Eubacterium tenue* | 5.801 | 10.948 | 3.972E-06 | 1.061E-05 |
| *Aestuariibacter halophilus* | 5.789 | 10.941 | 1.074E-05 | 2.399E-05 |
| *Lentibacter algarum* | 5.586 | 10.749 | 8.574E-06 | 1.968E-05 |
| *Clostridium thermosuccinogenes* | 5.576 | 10.732 | 2.431E-06 | 7.621E-06 |
| *Pseudoalteromonas haloplanktis* | 5.541 | 10.685 | 7.903E-08 | 6.134E-07 |
| *Mesorhizobium loti* | 5.431 | 10.593 | 8.378E-07 | 3.339E-06 |
| *Oceanirhabdus sediminicola* | 5.420 | 10.593 | 1.106E-05 | 2.403E-05 |
| *Thalassolituus oleivorans* | 5.405 | 10.553 | 3.108E-08 | 3.166E-07 |
| *Ketogulonicigenium vulgare* | 5.362 | 10.521 | 1.754E-07 | 1.077E-06 |
| *Tissierella creatinini* | 5.257 | 10.443 | 1.833E-05 | 3.727E-05 |
| *Vibrio vulnificus* | 5.169 | 10.334 | 2.574E-08 | 2.997E-07 |
| *Alkaliphilus peptidifermentans* | 5.054 | 10.241 | 5.114E-07 | 2.336E-06 |
| *Roseovarius aestuarii* | 4.988 | 10.185 | 9.409E-07 | 3.573E-06 |
| *Bythopirellula goksoyri* | 4.962 | 10.180 | 3.012E-05 | 5.644E-05 |
| *Tissierella praeacuta* | 4.791 | 10.023 | 2.896E-05 | 5.489E-05 |
| *Myroides indicus* | 4.649 | 9.901 | 4.199E-05 | 7.690E-05 |
| *Tindallia magadiensis* | 4.619 | 9.870 | 1.852E-05 | 3.727E-05 |
| *Marinifilum albidiflavum* | 4.582 | 9.837 | 1.589E-05 | 3.278E-05 |
| *Epibacterium multivorans* | 4.538 | 9.796 | 1.096E-05 | 2.403E-05 |
| *Rhizobium tropici* | 4.513 | 9.774 | 1.125E-05 | 2.412E-05 |
| *Cohaesibacter gelatinilyticus* | 4.399 | 9.684 | 4.486E-05 | 8.124E-05 |
| *Amphritea ceti* | 4.285 | 9.572 | 6.391E-06 | 1.551E-05 |
| *Hoeflea phototrophica* | 4.195 | 9.521 | 2.868E-04 | 4.583E-04 |
| *Cohaesibacter haloalkalitolerans* | 4.186 | 9.505 | 8.503E-05 | 1.444E-04 |
| *Pseudoteredinibacter isopora* | 4.144 | 9.473 | 1.133E-04 | 1.903E-04 |
| *Vibrio cholerae* | 4.140 | 9.464 | 5.581E-05 | 9.996E-05 |
| *Clostridium scindens* | 4.040 | 9.388 | 2.032E-04 | 3.280E-04 |
| *Sunxiuqinia elliptica* | 3.992 | 9.352 | 3.222E-04 | 5.098E-04 |
| *Streptomyces lazareus* | 3.931 | 9.297 | 1.627E-04 | 2.652E-04 |
| *Kordiimonas gwangyangensis* | 3.665 | 9.089 | 3.531E-04 | 5.533E-04 |

**Table 6.** Bacterial species enriched in clinically normal coral hosts as determined by edgeR contrast analysis. Designations are predicated at a false discovery rate (FDR) < 0.01. Shown are the logarithm of 2-fold change (log2FC), the logarithm of counts per million reads (log_2_CPM), the Benjamini-Hochberg (B-H) adjusted P value (Pvalue), and the FDR. The table is sorted from highest to lowest abundance (log_2_CPM).

| Bacterial Species | log2FC | Log2CPM | Pvalue | FDR |
| --- | --- | --- | --- | --- |
| *Heliobacterium oregonensis* | -10.362 | 15.430 | 6.492E-08 | 5.569E-07 |
| *Aureivirga marina* | -10.051 | 15.121 | 2.288E-07 | 1.243E-06 |
| *Blastocatella fastidiosa* | -3.996 | 14.774 | 5.076E-03 | 7.388E-03 |
| *Chloroflexus aurantiacus* | -3.283 | 13.728 | 4.550E-03 | 6.742E-03 |
| *Mycoplasma mycoides* | -4.034 | 13.702 | 1.188E-03 | 1.809E-03 |
| *Spirosoma navajo* | -8.304 | 13.383 | 2.275E-08 | 2.852E-07 |
| *Muricauda lutaonensis* | -7.559 | 12.652 | 9.644E-07 | 3.573E-06 |
| *Iamia majanohamensis* | -7.537 | 12.632 | 1.785E-07 | 1.077E-06 |
| *Streptococcus sanguinis* | -7.125 | 12.227 | 9.605E-07 | 3.573E-06 |
| *Candidatus Solibacter usitatus* | -6.976 | 12.083 | 3.459E-07 | 1.762E-06 |
| *Spirochaeta halophila* | -6.873 | 11.983 | 2.153E-07 | 1.210E-06 |
| *Bacillus licheniformis* | -6.848 | 11.958 | 1.708E-07 | 1.077E-06 |
| *Klebsiella pneumoniae* | -6.813 | 11.925 | 8.615E-09 | 1.560E-07 |
| *Vibrio* sp. Cl G9 | -6.470 | 11.595 | 3.172E-06 | 9.071E-06 |
| *Marinoscillum furvescens* | -6.464 | 11.591 | 7.350E-07 | 3.153E-06 |
| *Dictyoglomus turgidum* | -6.451 | 11.575 | 6.448E-08 | 5.569E-07 |
| *Candidatus Phytoplasma fraxini* | -6.442 | 11.567 | 5.657E-09 | 1.153E-07 |
| *Fulvivirga kasyanovii* | -6.435 | 11.563 | 1.025E-07 | 7.596E-07 |
| *Candidatus Liberibacter asiaticus* | -6.419 | 11.545 | 1.214E-08 | 1.979E-07 |
| *Aeromonas caviae* | -6.406 | 11.533 | 1.586E-07 | 1.077E-06 |
| *Aciditerrimonas ferrireducens* | -6.282 | 11.421 | 2.604E-06 | 8.009E-06 |
| *Arthrobacter davidanieli* | -6.194 | 11.331 | 6.928E-07 | 3.052E-06 |
| *Labilibacter aurantiacus* | -5.993 | 11.144 | 6.794E-06 | 1.605E-05 |
| *Tamlana agarivorans* | -5.991 | 11.140 | 1.557E-06 | 5.226E-06 |
| *Oenococcus oeni* | -5.984 | 11.133 | 7.049E-08 | 5.745E-07 |
| *Algisphaera agarilytica* | -5.923 | 11.079 | 8.398E-07 | 3.339E-06 |
| *Eionea nigra* | -5.829 | 10.990 | 6.472E-06 | 1.551E-05 |
| *Ilumatobacter fluminis* | -5.552 | 10.737 | 3.232E-06 | 9.082E-06 |
| *Sinorhizobium medicae* | -5.397 | 10.586 | 1.837E-10 | 1.497E-08 |
| *Pseudomonas marincola* | -5.360 | 10.558 | 9.923E-07 | 3.594E-06 |
| *Gloeobacter violaceus* | -5.269 | 10.472 | 2.584E-07 | 1.359E-06 |
| *Pseudomonas putida* | -5.245 | 10.452 | 1.275E-06 | 4.504E-06 |
| *Moraxella catarrhalis* | -5.109 | 10.326 | 7.350E-10 | 3.993E-08 |
| *Bartonella bacilliformis* | -5.068 | 10.290 | 1.052E-09 | 4.288E-08 |
| *Bdellovibrio bacteriovorus* | -4.794 | 10.052 | 1.997E-05 | 3.970E-05 |
| *Pectobacterium atrosepticum* | -4.780 | 10.040 | 1.274E-05 | 2.662E-05 |
| *Sulfobacillus thermosulfidooxidans* | -4.777 | 10.036 | 8.062E-07 | 3.339E-06 |
| *Serratia liquefaciens* | -4.732 | 9.998 | 3.617E-06 | 9.955E-06 |
| *Haloferula chungangensis* | -4.553 | 9.845 | 2.867E-05 | 5.489E-05 |
| *Roseivirga marina* | -4.509 | 9.807 | 5.499E-06 | 1.379E-05 |
| *Thiopropfundum lithotrophicum* | -4.437 | 9.743 | 1.379E-07 | 9.776E-07 |
| *Aridibacter kavangonensis* | -4.347 | 9.671 | 3.665E-06 | 9.955E-06 |
| *Coxiella burnetii* | -4.084 | 9.456 | 4.122E-05 | 7.635E-05 |
| *Gimesia maris* | -4.078 | 9.452 | 2.079E-05 | 4.084E-05 |
| *Thalassotalea agarivorans* | -4.006 | 9.393 | 6.572E-05 | 1.152E-04 |
| *Roseivirga spongicola* | -3.970 | 9.365 | 2.736E-05 | 5.309E-05 |
| *Lawsonella clevelandensis* | -3.882 | 9.296 | 8.244E-05 | 1.415E-04 |
| *Spiribacter curvatus* | -3.762 | 9.203 | 7.087E-05 | 1.229E-04 |
| *Ruegeria intermedia* | -3.647 | 9.113 | 5.797E-05 | 1.027E-04 |
| *Lewinella cohaerens* | -3.603 | 9.083 | 1.482E-04 | 2.441E-04 |
| *Ekhidna lutea* | -3.299 | 8.862 | 1.211E-04 | 2.015E-04 |
| *Gluconobacter oxydans* | -3.248 | 8.824 | 5.105E-04 | 7.849E-04 |
| *Portibacter lacus* | -3.118 | 8.736 | 3.608E-04 | 5.601E-04 |

**Table 7.** Bacterial species present in both diseased and clinically normal coral host samples. Designations are predicated at a false discovery rate (FDR) < 0.01. Shown are the logarithm of 2-fold change (log2FC), the logarithm of counts per million reads (log_2_CPM), the Benjamini-Hochberg (B-H) adjusted P value (Pvalue), and the false discovery rate (FDR). The table is sorted from highest to lowest abundance (log_2_CPM).

| Bacterial Species | log2FC | logCPM | Pvalue | FDR |
| --- | --- | --- | --- | --- |
| *Prosthecochloris vibrioformis* | -0.204 | 16.350 | 8.764E-01 | 8.929E-01 |
| "*Candidatus* Amoebophilus asiaticus" | -2.296 | 16.193 | 7.091E-02 | 9.713E-02 |
| *Arcobacter* sp. UDC415 | 2.027 | 15.337 | 1.499E-01 | 1.894E-01 |
| *Arcobacter nitrofigilis* | 1.707 | 14.929 | 3.458E-01 | 3.969E-01 |
| *Rhodospirillum rubrum* | -2.420 | 14.804 | 7.877E-02 | 1.061E-01 |
| *Achromobacter xylosoxidans* | -1.509 | 14.608 | 1.575E-01 | 1.975E-01 |
| *Aquimarina muelleri* | 1.041 | 13.905 | 5.184E-01 | 5.788E-01 |
| *Roseivirga seohaensis* | -0.235 | 13.847 | 8.593E-01 | 8.865E-01 |
| *Halodesulfobivibrio spirochaetisodalis* | 2.843 | 13.371 | 6.094E-02 | 8.563E-02 |
| *Tepidicaulis marinus* | 3.848 | 13.261 | 7.222E-03 | 1.042E-02 |
| *Candidatus* Pelagibacter ubique | -1.488 | 13.253 | 2.255E-01 | 2.723E-01 |
| *Fulvivirga imtechensis* | -1.281 | 12.590 | 2.811E-01 | 3.344E-01 |
| *Halodesulfobivibrio oceani* | -0.620 | 12.524 | 6.381E-01 | 6.843E-01 |
| *Halodesulfobivibrio marinisediminis* | -2.349 | 12.495 | 8.180E-02 | 1.093E-01 |
| *Achromobacter denitrificans* | -1.600 | 12.418 | 1.092E-01 | 1.435E-01 |
| *Pectobacterium carotovorum* | 0.404 | 12.300 | 7.613E-01 | 7.955E-01 |
| *Pirellula* sp. | 0.468 | 12.241 | 6.948E-01 | 7.402E-01 |
| *Pseudovibrio denitrificans* | 2.105 | 12.177 | 1.152E-01 | 1.467E-01 |
| *Aquisalinus flavus* | 1.354 | 12.177 | 3.023E-01 | 3.530E-01 |
| *Alteromonas macleodii* | -0.096 | 12.151 | 9.073E-01 | 9.073E-01 |
| *Wukongibacter baidiensis* | -0.132 | 12.088 | 9.050E-01 | 9.073E-01 |
| *Kiloniella spongiae* | 1.650 | 12.028 | 2.665E-01 | 3.194E-01 |
| *Clostridium botulinum* | -0.555 | 11.918 | 5.812E-01 | 6.428E-01 |
| *Nitrincola laciaponensis* | -1.234 | 11.911 | 1.692E-01 | 2.074E-01 |
| *Rubritalea tangerina* | 1.993 | 11.909 | 8.334E-02 | 1.104E-01 |
| *Azospirillum brasilense* | -2.278 | 11.752 | 4.922E-02 | 7.013E-02 |
| *Thalassobius mediterraneus* | 1.691 | 11.707 | 1.601E-01 | 1.987E-01 |
| *Achromobacter marplatensis* | -1.708 | 11.609 | 7.195E-02 | 9.774E-02 |
| *Pseudoalteromonas tetraodonis* | 1.576 | 11.576 | 4.948E-02 | 7.013E-02 |
| *Microcystis aeruginosa* | -0.183 | 11.557 | 8.340E-01 | 8.659E-01 |
| *Fulvivirga lutimaris* | -1.187 | 11.513 | 3.032E-01 | 3.530E-01 |
| *Sinorhizobium fredii* | -0.496 | 11.482 | 5.837E-01 | 6.428E-01 |
| *Fabibacter misakiensis* | -0.160 | 11.468 | 8.690E-01 | 8.908E-01 |
| *Riemerella anatipestifer* | 0.478 | 11.461 | 7.176E-01 | 7.595E-01 |
| *Prosthecochloris indica* | 0.906 | 11.447 | 4.719E-01 | 5.305E-01 |
| *Mesoplasma florum* | -1.018 | 11.403 | 3.309E-01 | 3.826E-01 |
| *Amphiplicatus metriotheophilus* | 0.592 | 11.367 | 6.276E-01 | 6.774E-01 |
| *Halodesulfobivrio aestuarii* | 1.369 | 11.365 | 3.006E-01 | 3.530E-01 |
| *Hoeflea halophila* | -0.880 | 11.325 | 4.411E-01 | 4.993E-01 |
| *Aeromonas salmonicida* | 0.162 | 11.313 | 8.865E-01 | 8.975E-01 |
| *Anaplasma platys* | -1.428 | 11.310 | 1.609E-01 | 1.987E-01 |
| *Phyllobacterium leguminum* | 2.050 | 11.304 | 1.137E-01 | 1.467E-01 |
| *Thermanaerovibrio acidaminovorans* | -2.019 | 11.214 | 6.707E-02 | 9.264E-02 |
| *Pelagibius litoralis* | -1.730 | 11.175 | 1.149E-01 | 1.467E-01 |
| *Mycoplasma gallisepticum* | -1.600 | 11.071 | 6.322E-02 | 8.808E-02 |
| *Woeseia oceani* | 0.591 | 11.061 | 6.016E-01 | 6.537E-01 |
| *Bartonella henselae* | -0.495 | 10.981 | 5.985E-01 | 6.537E-01 |
| *Lentisphaera profundii* | 1.572 | 10.951 | 1.150E-01 | 1.467E-01 |
| *Natranaerovirga hydrolytica* | 1.088 | 10.929 | 3.718E-01 | 4.238E-01 |
| *Acinetobacter radioresistens* | -0.387 | 10.845 | 7.250E-01 | 7.624E-01 |
| *Thermosynechococcus elongatus* | 1.385 | 10.806 | 1.772E-01 | 2.155E-01 |

### Inferred expression of major gene groupings

The inferred expression levels of the <u>Kyoto Encyclopedia of Genes and Genomes</u> (KEGG) major gene groupings including amino acid metabolism, biosynthesis of secondary metabolites, carbohydrate metabolism, energy metabolism, glycan biosynthesis and metabolism, lipid metabolism, metabolism of cofactors and vitamins, metabolism of amino acids, metabolism of terpenoids and polyketides, nucleotide metabolism, and xenobiotics biodegradation and metabolism were similar regardless of disease status and microbiome composition (FIG 4). Thus, major functionalities of the members of the clinically normal coral microbiomes are also present in equivalent numbers in the diseased coral microbiomes.

**FIG 4.**
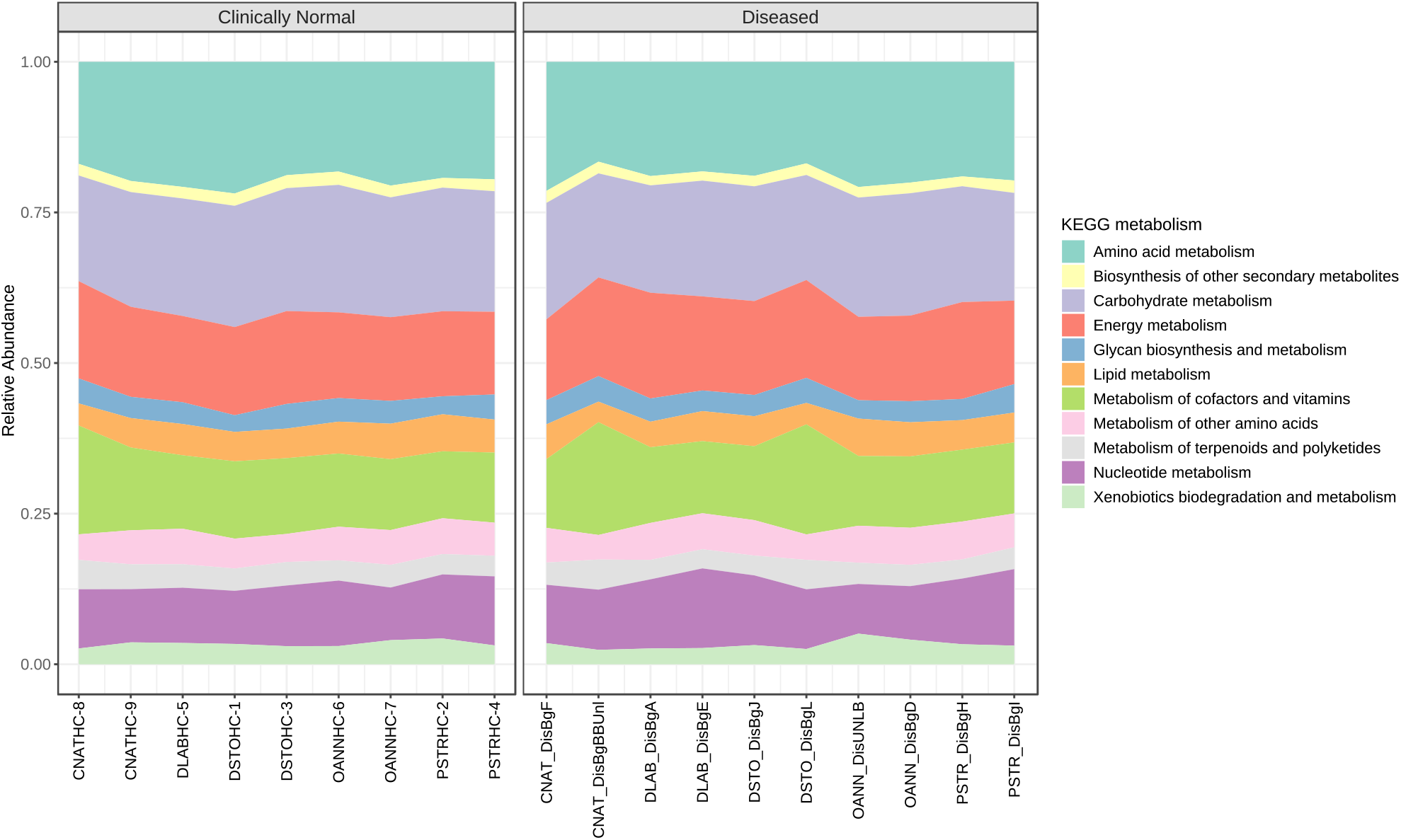
KEGG metabolism comparison of clinically normal and diseased coral microbiomes. Gene transcription was inferred using Tax4Fun.

## DISCUSSION

Stony Coral Tissue Loss Disease (SCTLD) is one of the more severe problems to confront Florida and Caribbean reefs in the 21^st^ century, potentially causing significant loss to ecosystem services and impacting economies. This outbreak is unusual due to its extended duration, apparent highly contagious nature, range of coral species affected, and rate of progression (https://floridakeys.noaa.gov/coral-disease/). Based on reports that amoxicillin can arrest tissue loss progression in several species, a number of researchers have focused on microbiome analyses to identify potential causative agents. J. L. Meyer et al. (25) recently reported identifying five amplicon sequence variants that were higher in abundance in diseased as opposed to healthy specimens of *Montastraea cavernosa*, *Orbicella faveolata*, *Diploria labyrinthiformis* and *Dicocoenia stokesii*. The identified amplicon sequence variants were found to be from an unclassified genus of Flavobacteriales, sequences identified as *Fusibacter* (Clostridiales), *Planktotalea* (Rhodobacterales), *Algicola* (Alteromonadales), and *Vibrio* (Vibrionales). The authors also identified opportunistic or saprophytic colonizers including *Epsilonbacteraeota, Patescibacteria, Clostridiales, Bacteroidetes*, and *Rhodobacterales* that were also found to be enriched in SCTLD disease lesions.

### Bacterial taxa found to be differentially elevated in abundance according to disease state

The taxonomic classifications we report here were performed at the species level using a closed reference database and a classification first approach, whereas those of J. L. Meyer et al. (25) and more recently those of S. M. Rosales et al. (26) were obtained using a cluster first approach. Although both strategies are widely used, each has pros and cons (27). Nevertheless, many of the findings of the two studies have general similarities regarding taxa identified as differentially elevated in numbers in diseased host corals. Sequence classifications that we obtained from the clinically normal coral samples were found to represent mostly bacterial species often found in coastal marine environments. In contrast, we found that several differentially abundant sequence classifications obtained from the diseased coral samples relate to bacteria with known pathogenic potential. Particularly notable are those that are common in specimens with active disease signs across the five coral species studied and that are relatively high in abundance. These include, but are not limited to, *Clostridioides difficile, Romboutsia lituseburensis, Arcobacter bivalviorum, Algicola bacteriolytica, Vibrio sp*. r24, *Shimia aquaeponti, Burkholderia gladioli, and Pseudoalteromonas haloplanktis* (Tables 4 and 5). In fact, *Clostridioides difficile, Arcobacter bivalviorum*, and *Algicola bacteriolytica* were found without exception in every diseased coral specimen. *Burkholderia gladioli*, usually a plant pathogen, produces several inhibitory and (or) toxic substances, among them gladiolin, bongkrek acid, enaxyloxin, and toxoflavin which can be fatal to humans. It is unknown if these toxins affect corals and (or) lower vertebrates. The presence of *Clostridioides difficile*, *Romboutsia lituseburensis* and other species often associated with soil, human, and animal gut consortia may implicate surface runoff and (or) fecal contamination as a contributing factor to SCTLD.

### Bacterial taxa found not to be differentially elevated in abundance according to disease state

In addition to the bacterial species found to be differentially elevated in abundance according to disease state, some prominent species’ numbers were found to be relatively constant regardless of disease state, and therefore their function(s) as microbiome members are important to consider. *Achromobacter xylosoxidans, Achromobacter denitrificans, Prosthecochloris vibrioformis,* and *“Candidatus* Amoebophilus asiaticus*”* were in this class as were several species of *Halodesulfovibrio*. *Achromobacter xylosoxidans* has been identified as an amoeba-resistant bacterium (ARB) in several studies (28, 29). Many free-living amoebas including *Acanthamoeba, Dictyostelium, Hartmannella*, and *Naegleria* can harbor bacteria (30, 31). ARBs have defense mechanisms that allow them to avoid ingestion or to survive in amoeba and other protists intracellularly and multiply. They may even persist through the resting cyst stage of the amoebae. Bacteria that can survive within amoebae gain protection from environmental insults from their amoeboid host (28, 32, 33). Because the survival mechanisms possessed by ARBs are similar to traits required for survival during infection of higher eukaryotes, ARBs are often found to be pathogenic (28, 29, 34–37).

*“Candidatus* Amoebophilus asiaticus” has been found widely in Caribbean corals (22) and was found to be among the dominant members of both the clinically normal and diseased Looe Key coral microbiomes sampled. Although it is an obligate endosymbiont of *Acanthamoeba* of the T4 grouping, it has not been possible to transfer it to other amoeba to date (38). This potentially indicates that this microbe is not widely amoeba-compatible or that its primary host may be something other than amoebae. Analysis of the genome of “*Ca.* Amoebophilus asiaticus” (39) revealed that it shares eukaryotic domains that are significantly enriched in the genomes of other intracellular parasitic bacteria (including chlamydiae*, Legionella pneumophila*, *Rickettsia bellii, Francisella tularensis, and Mycobacterium avium*). Thus, these diverse bacteria appear to exploit common mechanisms for interaction with their hosts. It is not clear if *“Ca.* Amoebophilus asiaticus” is internal to coral-associated free-living amoebae, or if the bacterium infects coral amoebocytes (40). If internal, the parasitic nature of this microbe might predispose corals to biotic and abiotic disease. If other amoeba-resistant bacteria that are present in high abundance (i.e. *Achromobacter sp.* as shown by amoeba co-culture) are also intracellular, the combination could be further debilitating. Amoebae are considered “training grounds” for bacterial pathogens and “*Ca.* Amoebophilus asiaticus” has an impressive number of insertion sequences in its genome (39) as does *Achromobacter* (41–43) making horizontal gene transfer and dissemination of virulence elements very possible in the confinement of intracellular space.

*Endozoicomonas,* another taxon that is known to be endosymbiotic, can be dominant in many coral and marine invertebrate species (44–47) achieving abundances as high as 85% (40, 44). *Endozoicomonas* is known to be capable of intracellular as well as free-living lifestyles, is sensitive to dysbiosis of the microbiome, and is diminished in intracellular numbers under unfavorable conditions. *Endozoicomonas spp.* are generally thought to be associated with healthy hosts (22, 48), but exhibit multiple relationships with their hosts, transitioning through symbiotic, mutualistic, and parasitic roles opportunistically (46). This endosymbiont has been thought to provide for metabolism of dimethylsulfoniopropionate (DMSP; (49–53) but genes for processing of DMSP were not identified in sequenced species (45, 46). Recently, however, K. Tandon et al. (54) have identified a dominant coral bacterium, *Endozoicomonas acroporae* that does metabolize DMSP. Other than DMSP processing, this endosymbiont is thought to provide benefit to the host including nutritional functions (55), antimicrobial functions (56), and functions associated with algal symbiont interactions (44, 57). Interestingly however, *Endozoicomonas* can also be a pathogen for fish (58). The finding of low *Endozoicomonas* abundances (less than 1.6%) in the bacterial consortia of multiple coral species at Looe Key regardless of disease state may be an indication that the “clinically normal” corals were stressed and (or) infected but not showing overt disease at the time of sampling. Disease and mortality continued unabated throughout 2018 as SCTLD moved southward along the Florida reef tract.

### Potential nutrient influence on affected coral microbiomes

Sewage contamination and surface runoff has long been recognized as problematic for coral health (5). V. N. Bednarz et al. (59) found that diazotroph-derived nitrogen assimilation by scleractinian corals was dependent on their metabolic status. More recently, T. Lachnit et al. (60) found that coral microbiome exposure to nutrient-rich conditions leads to dysbiosis and disease development. Another study has examined the impacts of land-based sources of pollution on the microbiota of Southeast Florida coral reefs and found that the major influence was from sewage outfalls adjacent to reef tracts, but that runoff from inlet outfalls was substantial as well (61). B. E. Lapointe et al. (62) have recently reported on a three-decade study of nitrogen enrichment at Looe Key and found a connection with reef decline. Nutrient levels can also modulate parasitic infections as shown by J. G. Klinges et al. (63) who have recently identified a marine invertebrate-associated Rickettsiales parasite that the authors found is encouraged by nutrient enrichment to overgrow and weaken and (or) kill the host coral cells it infests.

Excess nutrient levels that have been identified along the Florida Reef Tract may also drive another adverse process in stony Caribbean corals that may explain SCTLD pathology. *Prosthecochloris vibrioformis*, found in great abundance in all our coral specimens, is a strictly anaerobic green sulfur endolithic bacterium that has nitrogen assimilation and reduction pathways and produces proline in nitrogen rich environments (64). Proline has been found to be essential for the growth of *C. difficile* as well as other pathogens (65). *C. difficile* is a strict anaerobe that would be expected to locate in coral anoxic zones such as in the skeleton or at the skeletal-tissue interface. *C. difficile* produces toxins and is also known to produce para-cresol (p-cresol), a compound that is inhibitory to many Gram-negative bacteria that is thought to play a role in the ability of *C. difficile* to colonize tissues (66). An abundant species that we found ubiquitous in our coral specimens, *Achromobacter xylosoxidans,* can degrade p-cresol via p-cresol methylhydroxylase (67) unlike many other Gram-negative bacteria and would be immune to the effects of p-cresol. Additionally, although *Achromobacter xylosoxidans* is generally classified as aerobic, it can function anaerobically via denitrification (41). *Clostridia* have been found to be present in high abundance in coral skeleton and *Prosthecochloris vibrioformis* is dominant in stony corals due to the density of their skeletons providing a very anaerobic environment (64). Thus, it is possible that excess nutrients fuel *Prosthecochloris vibrioformis* growth and nitrogen fixation that subsequently supports *Clostridioides difficile* colonization, growth, and coral tissue damage via toxin production. The production of toxins at the tissue-skeletal interface may explain the observation that SCTLD pathology first affects the basal body wall of the coral polyps.

### Taxonomic redundancy

Finally, we tested to what degree members of the clinically normal and diseased coral microbiomes had similar inferred functionality profiles. Beta diversity analysis demonstrated that the microbes associated with the two disease states were quite different. The inferred expression of major gene groupings for these two groups, however, was similar regardless of disease status and microbiome composition. This result is similar to that found in other complex microbiomes including the human gut (13) and the algae, *Ulva* (17, 68) and reveals the redundancy of metabolic potential in coral-associated bacteria and the fluidity of specific bacterial species membership in the microbiomes of clinically normal and diseased stony corals.

### Conclusion

While disease outbreaks are not uncommon, this event is unique due to its large geographic range, extended duration, rapid progression, high rates of mortality and the number of species affected. The disease is thought to be caused by bacteria and can be transmitted to other corals through direct contact and water circulation (11). This study reveals the bacterial component of SCTLD specifically at a place and a time associated with a disease that is moving rapidly both geographically and temporally (69). Our inferences are made based on the abundance of bacterial sequences in diseased versus clinically normal states. Abundance is not an unequivocal measure of causality, however. It is possible that some rare, but influential bacterium somehow triggers remodeling of the coral microbiome (70, 71) given the variety of interactions among coral-associated bacteria (72). The redundancy of the microbiome members functional repertoire observed in this study as well as others (73) suggests that the makeup of the coral microbiome is a rather fluid thing influenced by space, time and abiotic factors. The dysbiosis generated from transient membership changes and microbiome remodeling in response to environmental circumstances may result in bacterial cliques being formed that collectively exhibit properties that have pathological implications for the host.

Unfortunately, it is not possible to pinpoint a specific causative agent for SCTLD at this juncture. It is possible, however, to offer some hypotheses regarding the SCTLD pathogenesis. It is tempting to conjecture, given the evidence presented here for example, that microbial endoparasites (such as “*Ca.* Amoebophilus asiaticus”) may predispose Caribbean corals to infection by opportunistic pathogens normally held at bay by a healthy microbiome including beneficial endosymbionts such as *Endozoicomonas* or that *Clostridioides difficile* introduced by terrestrial or surface runoff and fueled by excess nutrients may reside in the anoxic zone at the skeletal-tissue interface and produce the histopathological effects observed with SCTLD.

There may not be a single specific pathogen that causes SCTLD, but rather a polymicrobial mixture of bacteria each adding their “special touch” to the disease process and not necessarily being consistent in composition from one location to another or over time. We have identified a relatively short list of bacteria associated with SCTLD pathology. Our findings combined with those of prior and ongoing studies should help to guide the search for the causative agent(s) of this disease and their isolation into culture for detailed study. Proteomic and (or) metabolomic approaches may help to identify universal pathogenic mechanisms that operate regardless of the exact microbiome composition.

## MATERIALS AND METHODS

Five coral species (FIG 1) were targeted for collection on August 26, 2018 from two closely situated sites (separated by only 444 meters) at Looe Key, within the Florida Keys National Marine Sanctuary, (Table 1) under permit number FKNMS-2017-100, expiration date December 31, 2018. Duplicate specimens from actively diseased and clinically normal coral colonies were collected from *Orbicella annularis* (OANN), *Diploria labyrinthiformis* (DLAB), *Colpophyllia natans* (CNAT), *Pseudodiploria strigosa* (PSTR), and *Dichocoenia stokesii* (DSTO) except for *D. labyrinthiformis* for which we had only one clinically normal specimen (n=19 total specimens). Coral specimens were collected in the 2-5 meter depth range on scuba using hammer and clean chisels to collect fragments. Fragments were placed in zippered plastic bags for transport to the surface. Each of the coral specimens was placed into a separate 5-gal bucket filled with local natural seawater and covered with a lid. These samples were transported (approximately 1.5 h) to the Everglades National Park, Florida Bay Interagency Science Center (FBISC) in Key Largo, Florida for subsampling and processing. Samples were processed for metagenomic sequencing of bacterial suspensions from coral tissue homogenates. A subsample (approximately 3.25 cm^2^) that included tissue and skeleton from the disease margin of affected specimens or clinically normal specimens was aseptically removed and homogenized in a pre-sterilized mortar and pestle by grinding the tissue section in 2 mL of filter-sterilized artificial seawater (Instant Ocean Spectrum Brands, Blacksburg, VA) into a slurry. The homogenates were transferred into 2 mL sterile microcentrifuge tubes and centrifuged at 180 × *g* for 10 min to remove cellular debris. The supernatants (1 mL) containing bacteria were then recovered, bacteria were pelleted at 10,000 × g for 5 min, and the pellets were resuspended in 800 µL DNA/RNA Shield (Zymo Research, Irvine, CA) and frozen until DNA was extracted and used for direct 16S amplicon sequencing.

### DNA extraction, amplification, and sequencing

Genomic DNA from bacterial suspensions prepared from tissue homogenates was extracted and purified using the Zymo *Quick*-DNA Fecal/Soil Microbe Miniprep Kit (Zymo Research, Irvine, CA). All extracted DNA was stored in the supplied kit elution buffer at −20°C until polymerase chain reaction (PCR) was performed. Negative controls included kit reagent and field blanks. Positive controls were provided by extracts of a ZymoBIOMICS Microbial Community Standard (D6300; Zymo Research, Irvine, CA). Bacteria were characterized using PCR that targeted the V3 and V4 portion of the 16S rRNA gene to amplify that gene segment for subsequent Illumina MiSeq analysis. Amplicons were produced in two steps, first using rRNA-specific primers (see below) to generate a high concentration of input template followed by less efficient fusion primers that incorporate exogenous sequencing adapters. Oligonucleotide primers for the V3-V4 16S rRNA gene region (74, 75) were used to create a single amplicon of ∼460 base pairs (bp). The primers for the first amplification reaction were 16S Forward (5’ – CCTACGGGNGGCWGCAG – 3’) and 16S Reverse (5’ – GACTACHVGGGTATCTAATCC – 3’) (74, 76). Cycling parameters were: initial denaturation for 3 min at 95°C, followed by 35 cycles of 30 s at 95°C, 30 s at 55°C, and 30 s at 72°C, followed by a final extension at 72°C for 7 min. Amplicon size was confirmed by agarose gel electrophoresis using 5 µL of the reaction product. PCR products were cleaned with the Qiagen PCR Purification Kit (Valencia, CA) and quantified using a Qubit fluorometer (dsDNA HS Assay Kit, ThermoFisher Scientific, Grand Island, NY). Samples were diluted in 10 mM Tris buffer (pH 8.5) to a final concentration of 5 ng/µL. Using the 16S rRNA primers modified with the sequencing adaptors specified in Illumina’s 16S Metagenomic Sequencing Library Preparation (CT #: 15044223 Rev. B), amplicon libraries were prepared following the manufacturer’s protocol. Each sample was indexed with Illumina’s Nextera XT multiplex library indices, which incorporates two distinct 8 bp sequences on each end of the fragment. Libraries were quantified fluorometrically, as above. DNA size distribution was determined with an Agilent 2100 Bioanalyzer using the Agilent DNA 1000 Kit (Santa Clara, CA). The combined pool of indexed libraries was diluted to 4 nM using 10 mM Tris pH 8.5. A final 10 pM preparation was created with a 15% PhiX control spike and analyzed on a MiSeq 600 v3 cartridge. DNA sequences (16S) derived from the bacterial consortia of clinically normal and diseased coral samples as well as reagent and field blanks were treated identically.

### Sequence analysis

Sequences were classified taxonomically using the One Codex (https://www.onecodex.com/) pipeline (27, 77) by uploading machine-processed, quality filtered, and de-multiplexed FASTQ files to the One Codex web site for processing against the curated targeted database available there. Classification assignments were then downloaded from the One Codex web site as Microsoft Excel files. Low-abundance, non-informative classifications were removed from all data sets by trimming taxonomic assignments of sequencing reads from the bacterial component of each coral host to exclude those whose frequency was less than 0.25% of the total classification assignments associated with that coral host. Remaining classifications were then combined to obtain master lists of bacterial species found in clinically normal and diseased corals and ranked according to the number of reads classified to each member. Excel pivot tables were used to examine the distribution of identified bacterial species across coral species.

### Statistical analysis, diversity, and functional profiling inferred from bacterial taxonomy

Sequence classifications filtered as described above were also imported into MicrobiomeAnalyst (78) for further analyses including alpha and beta diversity to examine within and among sample diversity respectively, statistical analysis of bacterial association with disease state, and inference of biochemical pathways expressed in the consortia. Classifications additionally were filtered to remove those with a prevalence of less than 20% across samples and features with low variance (less than 10% based on the interquartile range). Data were transformed using the centered log ratio (CLR) method (79). Alpha diversity was calculated using the Simpson Index expressed as the compliment (ranging between zero and one where larger is more diverse) and tested for significance using Mann-Whitney or Kruskal-Wallis tests as appropriate. Beta diversity was examined using Analysis of Group Similarities (80) of Jenson-Shannon Divergence measures and visualized by NMDS (Nonmetric Multidimensional Scaling). Differences in the bacterial community composition between diseased and clinically normal corals were examined using the edgeR (81) method with an adjusted p value of 0.01. Finally, gene expression was inferred using the Tax4Fun module (82) of the MicrobiomeAnalyst package. The KEGG KO assignments that were generated were then assigned to major functional groupings and visualized with the MicrobiomeAnalyst Shotgun Data Profiling routines.

### Data availability

Raw sequences are available in the National Center for Biotechnology Information (NCBI) Sequence Read Archive (SRA) under project number PRJNA625928, accession numbers SAMN14596532 to SAMN14596550.

## ACKNOWLEDGEMENTS

Funding for this work was provided by the USGS Ecosystems Mission Area Fisheries Program and by NOAA’s Coral Reef Conservation Program Project # 1133 (C.M.W.).

We thank the National Park Service and the Florida Bay Interagency Science Center, Key Largo FL for providing laboratory and housing facilities and Christopher Kavanagh, Marine Ecologist for coordinating these arrangements and providing logistical support.

Roles in this research were equally shared by five scientists. W.B.S provided conceptualization, data curation, formal analysis, investigation, methodology, validation, visualization and writing the original draft manuscript. D.D.I. provided data curation, investigation, resources, writing – review and editing. K.M.B. provided funding acquisition, investigation, project administration, resources, supervision and writing – review and editing. C.M.W. provided funding acquisition, investigation, project administration, resources, and writing in review and editing. A.B. provided project administration and resources. K.N. provided resources and writing – review and editing.

Any use of trade, firm, or product names is for descriptive purposes only and does not imply endorsement by the U.S. Government.

Disclaimer: The views and analysis in this manuscript are solely those of the authors and do not necessarily reflect those of NOAA or National Ocean Service. The content of and findings within this document do not reflect NOAA policy.

## REFERENCES

1. Precht WF, Gintert BE, Robbart ML, Fura R, van Woesik R. 2016. Unprecedented Disease-Related Coral Mortality in Southeastern Florida. Sci Rep 6:31374.

2. FKNMS. 2020. Florida Reef Tract Coral Disease Outbreak: Disease, *on* Florida Keys National Marine Sanctuary, Office of National Marine Sanctuaries, National Oceanic and Atmospheric Administration, U.S. Department of Commerce. Accessed April 6.

3. Neely K. 2018. Surveying the Florida Keys Southern Coral Disease Boundary. Florida DEP, Miami, FL.

4. Pollock FJ, Morris PJ, Willis BL, Bourne DG. 2011. The Urgent Need for Robust Coral Disease Diagnostics. PLoS Path 7:e1002183.

5. Walker DI, Ormond RFG. 1982. Coral death from sewage and phosphate pollution at Aqaba, Red Sea. Mar Pollut Bull 13:21–25.

6. Neely KL. 2018. Ex-Situ Disease Treatment Trials. Florida DEP, Miami, FL.

7. Neely K, Macaulay K, Hower E, Dobler M. 2019. Effectiveness of topical antibiotics in treating corals affected by Stony Coral Tissue Loss Disease. bioRxiv 1:870402.

8. O’Neil K, Neely K, Patterson J. 2018. Nursery management of disease-raveged pillar coral (*Dendrogyra cylindrus*) on the Florida Reef Tract. Florida DEP, Miami, FL.

9. Voss J, Shilling E, Combs I. 2019. Intervention and fate tracking for corals affected by stony coral tissue loss disease in the northern Florida Reef Tract. Florida DEP, Miami, FL.

10. Walker B, Pitts K. 2019. Reef-building-coral Response to Amoxicillin Intervention and Broader-scale Coral Disease Intervention. Florida DEP, Miami, FL.

11. Aeby GS, Ushijima B, Campbell JE, Jones S, Williams GJ, Meyer JL, Häse C, Paul VJ. 2019. Pathogenesis of a Tissue Loss Disease Affecting Multiple Species of Corals Along the Florida Reef Tract. Frontiers in Marine Science 6.

12. Ahmad AF, Dwivedi G, O’Gara F, Caparros-Martin J, Ward NC. 2019. The gut microbiome and cardiovascular disease: current knowledge and clinical potential. American Journal of Physiology-Heart and Circulatory Physiology 317:H923–H938.

13. Huttenhower C, Gevers D, Knight R, Abubucker S, Badger JH, Chinwalla AT, Creasy HH, Earl AM, FitzGerald MG, Fulton RS, Giglio MG, Hallsworth-Pepin K, Lobos EA, Madupu R, Magrini V, Martin JC, Mitreva M, Muzny DM, Sodergren EJ, Versalovic J, Wollam AM, Worley KC, Wortman JR, Young SK, Zeng Q, Aagaard KM, Abolude OO, Allen-Vercoe E, Alm EJ, Alvarado L, Andersen GL, Anderson S, Appelbaum E, Arachchi HM, Armitage G, Arze CA, Ayvaz T, Baker CC, Begg L, Belachew T, Bhonagiri V, Bihan M, Blaser MJ, Bloom T, Bonazzi V, Paul Brooks J, Buck GA, Buhay CJ, Busam DA, Campbell JL, et al. 2012. Structure, function and diversity of the healthy human microbiome. Nature 486:207–214.

14. Segata N, Izard J, Waldron L, Gevers D, Miropolsky L, Garrett WS, Huttenhower C. 2011. Metagenomic biomarker discovery and explanation. Genome Biol 12:R60.

15. Verster AJ, Borenstein E. 2018. Competitive lottery-based assembly of selected clades in the human gut microbiome. Microbiome 6:186–186.

16. Bass D, Stentiford GD, Wang HC, Koskella B, Tyler CR. 2019. The Pathobiome in Animal and Plant Diseases. Trends Ecol Evol 34:996–1008.

17. Burke C, Thomas T, Lewis M, Steinberg P, Kjelleberg S. 2011. Composition, uniqueness and variability of the epiphytic bacterial community of the green alga Ulva australis. ISME J 5:590–600.

18. Elshahed MS, Najar FZ, Aycock M, Qu C, Roe BA, Krumholz LR. 2005. Metagenomic analysis of the microbial community at Zodletone Spring (Oklahoma): insights into the genome of a member of the novel candidate division OD1. Appl Environ Microbiol 71:7598–602.

19. Galbraith H, Iwanowicz D, Spooner D, Iwanowicz L, Keller D, Zelanko P, Adams C. 2018. Exposure to Synthetic Hydraulic Fracturing Waste Influences the Mucosal Bacterial Community Structure of the Brook Trout (*Salvelinus fontinalis*) Epidermis. AIMS Microbiology 4:413–427.

20. Ainsworth T, Krause L, Bridge T, Torda G, Raina J-B, Zakrzewski M, Gates RD, Padilla-Gamiño JL, Spalding HL, Smith C, Woolsey ES, Bourne DG, Bongaerts P, Hoegh-Guldberg O, Leggat W. 2015. The coral core microbiome identifies rare bacterial taxa as ubiquitous endosymbionts. The ISME Journal 9:2261–2274.

21. Bernasconi R, Stat M, Koenders A, Paparini A, Bunce M, Huggett MJ. 2019. Establishment of Coral-Bacteria Symbioses Reveal Changes in the Core Bacterial Community With Host Ontogeny. Front Microbiol 10.

22. Huggett MJ, Apprill A. 2019. Coral microbiome database: Integration of sequences reveals high diversity and relatedness of coral-associated microbes. Environ Microbiol Rep 11:372–385.

23. Pollock FJ, McMinds R, Smith S, Bourne DG, Willis BL, Medina M, Thurber RV, Zaneveld JR. 2018. Coral-associated bacteria demonstrate phylosymbiosis and cophylogeny. Nature Communications 9:4921.

24. Sweet MJ, Brown BE, Dunne RP, Singleton I, Bulling M. 2017. Evidence for rapid, tide-related shifts in the microbiome of the coral *Coelastrea aspera*. Coral Reefs 36:815–828.

25. Meyer JL, Castellanos-Gell J, Aeby GS, Häse CC, Ushijima B, Paul VJ. 2019. Microbial Community Shifts Associated With the Ongoing Stony Coral Tissue Loss Disease Outbreak on the Florida Reef Tract. Front Microbiol 10.

26. Rosales SM, Clark AS, Huebner LK, Ruzicka RR, Muller EM. 2020. *Rhodobacterales* and *Rhizobiales* Are Associated With Stony Coral Tissue Loss Disease and Its Suspected Sources of Transmission. Front Microbiol 11.

27. Siegwald L, Touzet H, Lemoine Y, Hot D, Audebert C, Caboche S. 2017. Assessment of Common and Emerging Bioinformatics Pipelines for Targeted Metagenomics. PLoS One 12:e0169563.

28. Greub G, La Scola B, Raoult D. 2004. Amoebae-resisting bacteria isolated from human nasal swabs by amoebal coculture. Emerg Infect Dis 10:470–7.

29. Thomas V, McDonnell G, Denyer SP, Maillard J-Y. 2010. Free-living amoebae and their intracellular pathogenic microorganisms: risks for water quality. FEMS Microbiol Rev 34:231–259.

30. Strassmann JE, Shu L. 2017. Ancient bacteria–amoeba relationships and pathogenic animal bacteria. PLoS Biol 15:e2002460.

31. Denoncourt AMVEPSJC. 2014. Potential Role of Bacteria Packaging by Protozoa in the Persistence and Transmission of Pathogenic Bacteria. Front Microbiol 5:240.

32. Guimaraes AJ, Gomes KX, Cortines JR, Peralta JM, Peralta RH. 2016. *Acanthamoeba spp.* as a universal host for pathogenic microorganisms: One bridge from environment to host virulence. Microbiol Res 193:30–38.

33. Hilbi H, Weber SS, Ragaz C, Nyfeler Y, Urwyler S. 2007. Environmental predators as models for bacterial pathogenesis. Environ Microbiol 9:563–75.

34. Essig A, Heinemann M, Simnacher U, Marre R. 1997. Infection of *Acanthamoeba castellanii* by *Chlamydia pneumoniae*. Appl Environ Microbiol 63:1396–9.

35. Jacquier N, Aeby, S., Lienar, J., Greub, G. 2013. Discovery of New Intracellular Pathogens by Amoebal Coculture and Amoebal Enrichment Approaches. Journal of Visualized Experiments 80:51055.

36. Janda WM. 2010. Amoeba-Resistant Bacteria: Their Role in Human Infections. Clin Microbiol Newsl 32:177–184.

37. Long JJ, Jahn CE, Sanchez-Hidalgo A, Wheat W, Jackson M, Gonzalez-Juarrero M, Leach JE. 2018. Interactions of free-living amoebae with rice bacterial pathogens *Xanthomonas oryzae* pathovars oryzae and oryzicola. PLoS One 13:e0202941.

38. Horn M, Harzenetter MD, Linner T, Schmid EN, Muller KD, Michel R, Wagner M. 2001. Members of the *Cytophaga-Flavobacterium-Bacteroides* phylum as intracellular bacteria of acanthamoebae: proposal of ‘*Candidatus* Amoebophilus asiaticus’. Environ Microbiol 3:440–9.

39. Schmitz-Esser S, Tischler P, Arnold R, Montanaro J, Wagner M, Rattei T, Horn M. 2010. The genome of the amoeba symbiont “*Candidatus* Amoebophilus asiaticus” reveals common mechanisms for host cell interaction among amoeba-associated bacteria. J Bacteriol 192:1045–57.

40. Apprill A, Weber LG, Santoro AE. 2016. Distinguishing between Microbial Habitats Unravels Ecological Complexity in Coral Microbiomes. mSystems 1:e00143–16.

41. Jakobsen TH, Hansen MA, Jensen PØ, Hansen L, Riber L, Cockburn A, Kolpen M, Rønne Hansen C, Ridderberg W, Eickhardt S, Hansen M, Kerpedjiev P, Alhede M, Qvortrup K, Burmølle M, Moser C, Kühl M, Ciofu O, Givskov M, Sørensen SJ, Høiby N, Bjarnsholt T. 2013. Complete Genome Sequence of the Cystic Fibrosis Pathogen *Achromobacter xylosoxidans* NH44784-1996 Complies with Important Pathogenic Phenotypes. PLoS One 8:e68484.

42. Jeukens J, Freschi L, Vincent AT, Emond-Rheault JG, Kukavica-Ibrulj I, Charette SJ, Levesque RC. 2017. A pan-genomic approach to understand the basis of host adaptation in *Achromobacter*. Genome Biol Evol 9:1030–1046.

43. Traglia GM, Almuzara M, Merkier AK, Adams C, Galanternik L, Vay C, Centrón D, Ramírez MS. 2012. *Achromobacter xylosoxidans*: An Emerging Pathogen Carrying Different Elements Involved in Horizontal Genetic Transfer. Curr Microbiol 65:673–678.

44. Morrow KM, Moss AG, Chadwick NE, Liles MR. 2012. Bacterial associates of two Caribbean coral species reveal species-specific distribution and geographic variability. Appl Environ Microbiol 78:6438–6449.

45. Neave MJ, Michell CT, Apprill A, Voolstra CR. 2014. Whole-genome sequences of three symbiotic *Endozoicomonas* strains. Genome announcements 2:e00802–14.

46. Neave MJ, Michell CT, Apprill A, Voolstra CR. 2017. *Endozoicomonas* genomes reveal functional adaptation and plasticity in bacterial strains symbiotically associated with diverse marine hosts. Sci Rep 7:40579.

47. Schill WB, Iwanowicz D, Adams C. 2017. *Endozoicomonas* Dominates the Gill and Intestinal Content Microbiomes of *Mytilus edulis* from Barnegat Bay, New Jersey. J Shellfish Res 36:391–401, 11.

48. Ding J-Y, Shiu J-H, Chen W-M, Chiang Y-R, Tang S-L. 2016. Genomic Insight into the Host– Endosymbiont Relationship of *Endozoicomonas montiporae* CL-33T with its Coral Host. Front Microbiol 7.

49. Raina JB, Clode PL, Cheong S, Bougoure J, Kilburn MR, Reeder A, Foret S, Stat M, Beltran V, Thomas-Hall P, Tapiolas D, Motti CM, Gong B, Pernice M, Marjo CE, Seymour JR, Willis BL, Bourne DG. 2017. Subcellular tracking reveals the location of dimethylsulfoniopropionate in microalgae and visualises its uptake by marine bacteria. Elife 6.

50. Raina JB, Dinsdale EA, Willis BL, Bourne DG. 2010. Do the organic sulfur compounds DMSP and DMS drive coral microbial associations? Trends Microbiol 18:101–8.

51. Raina JB, Tapiolas D, Motti CA, Foret S, Seemann T, Tebben J, Willis BL, Bourne DG. 2016. Isolation of an antimicrobial compound produced by bacteria associated with reef-building corals. PeerJ 4:e2275.

52. Raina JB, Tapiolas D, Willis BL, Bourne DG. 2009. Coral-associated bacteria and their role in the biogeochemical cycling of sulfur. Appl Environ Microbiol 75:3492–501.

53. Raina JB, Tapiolas DM, Foret S, Lutz A, Abrego D, Ceh J, Seneca FO, Clode PL, Bourne DG, Willis BL, Motti CA. 2013. DMSP biosynthesis by an animal and its role in coral thermal stress response. Nature 502:677–80.

54. Tandon K, Chiang P-W, Lu C-Y, Yang S-H, Chen Y-F, Wada N, Chen P-Y, Chang H-Y, Chou M-S, Chen W-M, Tang S-L. 2019. Dominant coral bacterium *Endozoicomonas acroporae* metabolizes DMSP. bioRxiv doi:10.1101/519546:519546.

55. La Rivière M, Roumagnac M, Garrabou J, Bally M. 2013. Transient shifts in bacterial communities associated with the temperate gorgonian *Paramuricea clavata* in the Northwestern Mediterranean Sea. PLoS One 8:e57385–e57385.

56. Bourne D, Iida Y, Uthicke S, Smith-Keune C. 2008. Changes in coral-associated microbial communities during a bleaching event. ISME J 2:350–63.

57. Pantos O, Bongaerts P, Dennis PG, Tyson GW, Hoegh-Guldberg O. 2015. Habitat-specific environmental conditions primarily control the microbiomes of the coral *Seriatopora hystrix*. The ISME journal 9:1916–1927.

58. Mendoza M, Güiza L, Martinez X, Caraballo X, Rojas J, Aranguren LF, Salazar M. 2013. A novel agent (*Endozoicomonas elysicola*) responsible for epitheliocystis in cobia *Rachycentrum canadum* larvae. Dis Aquat Org 106:31–37.

59. Bednarz VN, Grover R, Maguer JF, Fine M, Ferrier-Pages C. 2017. The Assimilation of Diazotroph-Derived Nitrogen by Scleractinian Corals Depends on Their Metabolic Status. mBio 8.

60. Lachnit T, Bosch TCG, Deines P. 2019. Exposure of the Host-Associated Microbiome to Nutrient-Rich Conditions May Lead to Dysbiosis and Disease Development-an Evolutionary Perspective. mBio 10.

61. Staley C, Kaiser T, Gidley ML, Enochs IC, Jones PR, Goodwin KD, Sinigalliano CD, Sadowsky MJ, Chun CL. 2017. Differential Impacts of Land-Based Sources of Pollution on the Microbiota of Southeast Florida Coral Reefs. Appl Environ Microbiol 83:e03378–16.

62. Lapointe BE, Brewton RA, Herren LW, Porter JW, Hu C. 2019. Nitrogen enrichment, altered stoichiometry, and coral reef decline at Looe Key, Florida Keys, USA: a 3-decade study. Mar Biol 166:108.

63. Klinges JG, Rosales SM, McMinds R, Shaver EC, Shantz AA, Peters EC, Eitel M, Wörheide G, Sharp KH, Burkepile DE, Silliman BR, Vega Thurbe. RL. 2019. Phylogenetic, genomic, and biogeographic characterization of a novel and ubiquitous marine invertebrate-associated *Rickettsiales* parasite, “*Candidatus* Aquarickettsia rohweri”, gen. nov., sp. nov. The ISME Journal 13:2938–2953.

64. Yang SH, Tandon K, Lu CY, Wada N, Shih CJ, Hsiao SS, Jane WN, Lee TC, Yang CM, Liu CT, Denis V, Wu YT, Wang LT, Huang L, Lee DC, Wu YW, Yamashiro H, Tang SL. 2019. Metagenomic, phylogenetic, and functional characterization of predominant endolithic green sulfur bacteria in the coral *Isopora palifera*. Microbiome 7:3.

65. Christgen SL, Becker DF. 2017. Role of Proline in Pathogen and Host Interactions. Antioxidants & Redox Signaling 30:683–709.

66. Passmore IJ, Letertre MPM, Preston MD, Bianconi I, Harrison MA, Nasher F, Kaur H, Hong HA, Baines SD, Cutting SM, Swann JR, Wren BW, Dawson LF. 2018. Para-cresol production by *Clostridium difficile* affects microbial diversity and membrane integrity of Gram-negative bacteria. PLoS Path 14:e1007191–e1007191.

67. Hopper DJ, Bossert ID, Rhodes-Roberts ME. 1991. p-cresol methylhydroxylase from a denitrifying bacterium involved in anaerobic degradation of p-cresol. J Bacteriol 173:1298–1301.

68. Burke C, Steinberg P, Rusch D, Kjelleberg S, Thomas T. 2011. Bacterial community assembly based on functional genes rather than species. Proc Natl Acad Sci U S A 108:14288–93.

69. Muller EM, Sartor C, Alcaraz NI, van Woesik. R. 2020. Spatial Epidemiology of the Stony-Coral-Tissue-Loss Disease in Florida. Frontiers in Marine Science 7.

70. Banerjee S, Schlaeppi K, van der Heijden MGA. 2018. Keystone taxa as drivers of microbiome structure and functioning. Nature Reviews Microbiology 16:567–576.

71. Hajishengallis G, Darveau RP, Curtis MA. 2012. The keystone-pathogen hypothesis. Nature Reviews Microbiology 10:717–725.

72. Rypien KL, Ward JR, Azam F. 2010. Antagonistic interactions among coral-associated bacteria. Environ Microbiol 12:28–39.

73. Hernandez-Agreda A, Leggat W, Bongaerts P, Herrera C, Ainsworth TD. 2018. Rethinking the Coral Microbiome: Simplicity Exists within a Diverse Microbial Biosphere. mBio 9:e00812–18.

74. Herlemann DPR, Labrenz M, Jürgens K, Bertilsson S, Waniek JJ, Andersson AF. 2011. Transitions in bacterial communities along the 2000 km salinity gradient of the Baltic Sea. The ISME Journal 5:1571–1579.

75. Klindworth A, Pruesse E, Schweer T, Peplies J, Quast C, Horn M, Glöckner FO. 2012. Evaluation of general 16S ribosomal RNA gene PCR primers for classical and next-generation sequencing-based diversity studies. Nucleic Acids Res 41:e1–e1.

76. Klindworth A, Pruesse E, Schweer T, Peplies J, Quast C, Horn M, Glöckner FO. 2013. Evaluation of general 16S ribosomal RNA gene PCR primers for classical and next-generation sequencing-based diversity studies. Nucleic Acids Res 41:e1.

77. Minot SK, N.; Greenfield, N.B. 2015. One Codex: A sensitive and accurate data platform for genomic microbial identification. BioRxiv doi:http://dx.doiorg/10.1101/027607.

78. Dhariwal A, Chong J, Habib S, King IL, Agellon LB, Xia J. 2017. MicrobiomeAnalyst: a web-based tool for comprehensive statistical, visual and meta-analysis of microbiome data. Nucleic Acids Res 45:W180–W188.

79. Gloor GB, Wu JR, Pawlowsky-Glahn V, Egozcue JJ. 2016. It’s all relative: analyzing microbiome data as compositions. Ann Epidemiol 26:322–329.

80. Clarke KR. 1993. Non-parametric multivariate analyses of changes in community structure. Aust J Ecol 18:117–143.

81. Robinson MD, McCarthy DJ, Smyth GK. 2010. edgeR: a Bioconductor package for differential expression analysis of digital gene expression data. Bioinformatics (Oxford, England) 26:139–140.

82. Aßhauer KP, Wemheuer B, Daniel R, Meinicke P. 2015. Tax4Fun: predicting functional profiles from metagenomic 16S rRNA data. Bioinformatics 31:2882–2884.

